# Multistability and consequent phenotypic plasticity in AMPK-Akt double negative feedback loop in cancer cells

**DOI:** 10.1101/2020.12.21.423274

**Authors:** Adithya Chedere, Kishore Hari, Saurav Kumar, Annapoorni Rangarajan, Mohit Kumar Jolly

**Author notes:** These authors contributed equally. Authors to whom correspondence should be addressed (A.R.), (M.K.J.).

## Abstract

Adaptation and survival of cancer cells to various stress and growth factor conditions is crucial for successful metastasis. A double-negative feedback loop between two serine/threonine kinases AMPK and Akt can regulate the adaptation of breast cancer cells to matrix-deprivation stress. This feedback loop can generate majorly two phenotypes or cell states: matrix detachment-triggered pAMPK^high^/ pAkt^low^ state, and matrix (re)attachment-triggered pAkt^high^/ pAMPK^low^ state. However, whether these two cell states can exhibit phenotypic plasticity and heterogeneity in a given cell population, i.e., whether they can co-exist and undergo spontaneous switching to generate the other subpopulation, remains unclear. Here, we develop a mechanism-based mathematical model that captures the set of experimentally reported interactions among AMPK and Akt. Our simulations suggest that the AMPK-Akt feedback loop can give rise to two co-existing phenotypes (pAkt^high^/ pAMPK^low^ and pAMPK^high^/pAkt^low^) in specific parameter regimes. Next, to test the model predictions, we segregated these two subpopulations in MDA-MB-231 cells and observed that each of them was capable of switching to another in adherent conditions. Finally, the predicted trends are supported by clinical data analysis of TCGA breast cancer and pan-cancer cohorts that revealed negatively correlated pAMPK and pAkt protein levels. Overall, our integrated computational-experimental approach unravels that AMPK-Akt feedback loop can generate multistability and drive phenotypic switching and heterogeneity in a cancer cell population.

## Introduction

Despite major advances in cancer research, metastasis remains clinically unsolved and claims the vast majority of cancer-related deaths. Metastasis is a highly inefficient process with extremely high (>99.8%) rates of attrition. A hallmark of cancer cells that can successfully metastasize is their ability to dynamically adapt to their changing microenvironments, called as phenotypic plasticity or switching [1,2]. Thus, understanding the rules of phenotypic plasticity and identifying therapeutic perturbations to reduce the fitness of metastasizing cells can be crucial for restricting the disease aggressiveness. Metastasizing cells can often display phenotypic plasticity along multiple interconnected axes. Two most well-characterized axes are epithelial-mesenchymal plasticity (EMP) and cancer stem cell (CSC) plasticity [3,4]. More recently, the axes of metabolic plasticity (i.e., switching between more glycolytic vs. more oxidative phosphorylating states) [5,6] and drug resistance (i.e., switching between drugresistant and drug-sensitive states) [7,8] are being investigated. A hallmark of networks involved in plasticity along these axes is the presence of double negative or mutually inhibitory feedback loops that can generate two (or more) phenotypes that cells can acquire. In the context of EMP, ZEB1 forms such loops with miR-200 and GRHL2, thus giving rise to multiple states – epithelial (high miR-200 and GRHL2, low ZEB), mesenchymal (low miR-200 and GRHL2, high ZEB), and hybrid epithelial/ mesenchymal (medium levels of miR-200, GRHL2 and ZEB) [9]. Similarly, for CSC plasticity, LIN28 and let-7 inhibit each other [10], and for metabolic plasticity, HIF-1α and AMPK can inhibit each other [5]. The emergent dynamics of abovementioned loops has been thoroughly investigated, and they have been shown as capable of exhibiting multistability (i.e. co-existence of multiple phenotypes). Such multistability has been posited to underlie phenotypic switching; disrupting such loops can restrict phenotypic switching, as witnessed for feedback loops of ZEB1 with GRHL2 and miR-200 [11,12].

Recent work from our laboratory has uncovered a double-negative feedback loop between two serine/threonine kinases AMPK and Akt operating in adaptation of breast cancer cells to matrix-deprivation [13]. In epithelial cells, matrix-deprivation usually drives programmed cell death known as ‘anoikis’ [14], but few detached epithelial cells can develop resistance to anoikis [15,16]. AMPK (AMP-activated protein kinase) is activated in cells facing bioenergetic or metabolic stress and can switch on energy-generating catabolic processes such as glycolysis and inhibit energy-consuming anabolic processes such as lipid and protein synthesis [17]. Conversely, upon growth factor stimulation, Akt gets activated promoting anabolic processes of lipid and protein synthesis, driving cell growth and proliferation [18]. Upon matrix deprivation, AMPK is activated that drives upregulation of PHLPP2 protein levels which can inactivate Akt. On the other hand, upon matrix (re)attachment, Akt is activated which can repress AMPK activity through PP2Cα [13]. Thus, while adherent cells showed a (re)attachment-triggered pAkt^high^/ pAMPK^low^ state, matrix-deprived cells demonstrated a detachment-triggered pAMPK^high^/pAkt^low^ state. However, because this analysis was done at a population (or bulk) level, two questions remain to be answered: a) can these cell states/phenotypes co-exist in the same cell population?, and b) can these two subpopulations ‘spontaneously’ switch between themselves to give rise to one another?

Here, we adopt an integrative computational-experimental approach to answer these questions. First, we develop a mechanism-based mathematical model that captures the set of experimentally reported interactions among AMPK, Akt, PHLPP2 and PP2Cα. Simulations reveal that AMPK-Akt feedback loop can give rise to two phenotypes - pAkt^high^/ pAMPK^low^, and pAMPK^high^/pAkt^low^ - that can co-exist in specific parameter regimes, and switch between one another under the influence of biological noise.

Next, we segregated the two subpopulations in MDA-MB-231 cells and observed that under adherent conditions, each of them was capable of giving rise to another, thus validating our model prediction. Finally, clinical data analysis revealed a negative correlation between pAMPK and pAkt protein levels in TCGA breast cancer and pan-cancer cohorts. Overall, our results suggest that AMPK-Akt feedback loop can be bistable, and therefore drive phenotypic switching and non-genetic heterogeneity in a cancer cell population.

## Results

### AMPK-Akt feedback loop can give rise to two states: pAkt^high^/ pAMPK^low^ and pAMPK^high^/pAkt^low^

First, we gathered experimentally curated information about interconnections among AMPK and Akt.

AMPK and Akt can antagonistically regulate common downstream effectors such as mTOR signaling and FOXO signaling through differential phosphorylation [19]. Moreover, they can affect the activation status of one another. For instance, AMPK activating agents such as AICAR and phenformin can reduce the phosphorylation of Akt [20]. Adiponectin-activated AMPK can dephosphorylate Akt by increasing the activity of protein phosphatase 2A (PP2A) through dephosphorylating PP2Ac at Tyr307 [21]. On the other hand, upon insulin treatment, Akt is activated and it phosphorylates Ser487 of the serine/threonine rich loop (ST loop) in AMPK-α1 subunit, thus reducing subsequent phosphorylation and LKB1-or CAMKKβ-dependent AMPK activation at Thr172. Also, GSK3, another substrate of Akt, can phosphorylate the AMPK-α1 subunit at Thr481 and Ser477, further inhibiting AMPK activation by Thr172 phosphorylation [22]. Thus, AMPK and Akt pathways seem to inhibit the activity of one another.

Such a double negative association between AMPK and Akt was also reported in breast cancer cells during matrix attachment and detachment. The activation of AMPK upon matrix-deprivation drove upregulation of PHLPP2 protein levels, which can inactivate Akt. On the other hand, when cells were (re)attached to the matrix, Akt was activated that repressed AMPK activity through PP2Cα [13] (**Fig 1**). Put together, the abovementioned interactions reveal a mutually inhibitory feedback loop between the AMPK and Akt. This loop is reminiscent of “toggle switches” formed by various ‘master regulators’ of two (or more) diverse cell states, seen during embryonic development and disease progression [23]. Such feedback loops can occur at transcriptional [24,25], post-transcriptional [26,27], and cellcell communication levels [28,29]. Here, a ‘toggle switch’ is observed between two kinases.

**Figure 1:**
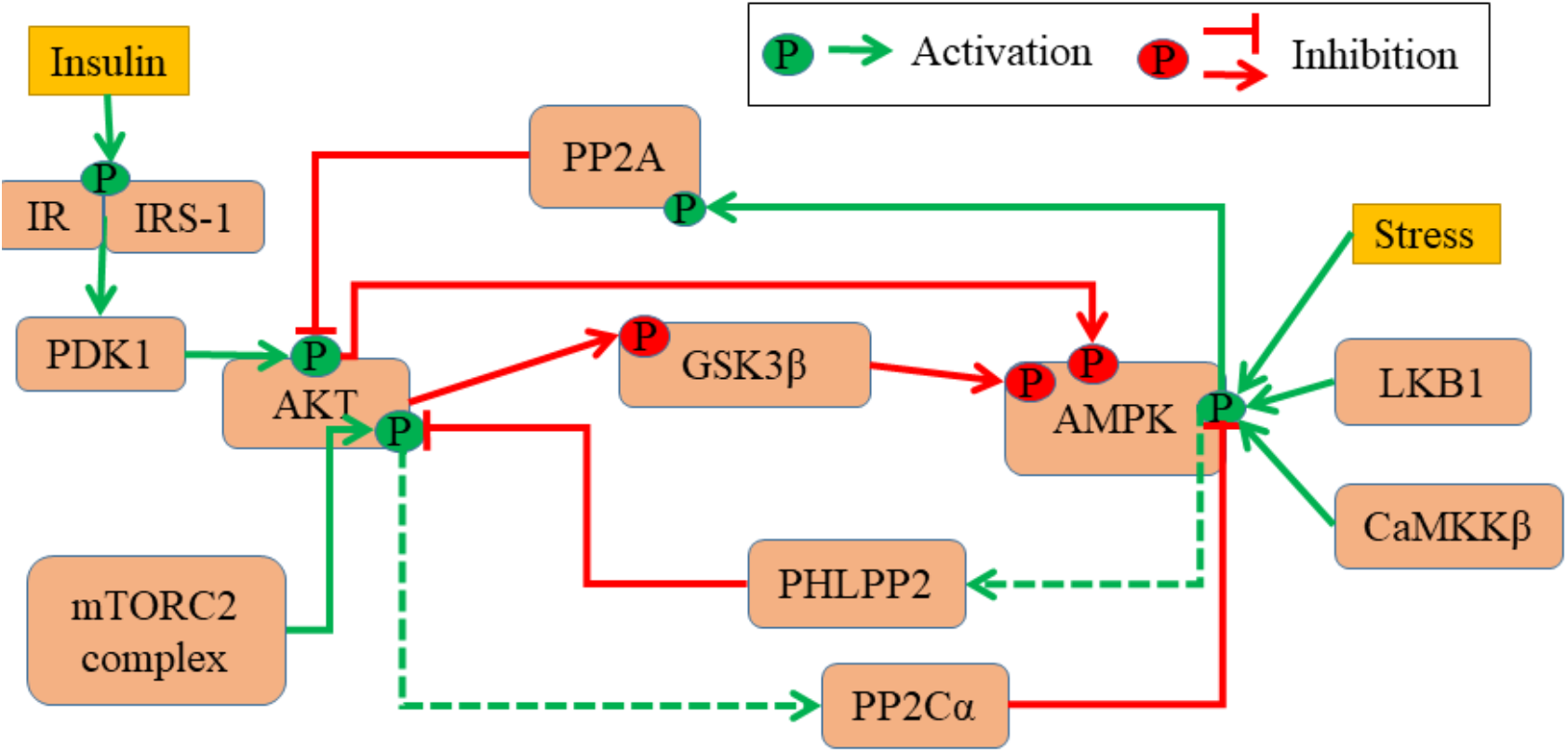
AMPK-Akt feedback loop. Regulatory network between AMPK and Akt. Green arrows and associated green ‘P’ circles denote activation by phosphorylation. Red arrows indicate deactivation by phosphorylation, and red hammerheads show deactivation by dephosphorylation. Solid lines show known molecular mechanisms, dashed lines show scenarios where such information is not available.

Next, we investigated the emergent dynamics of the AMPK-Akt feedback loop. For sake of simplicity, we considered the set of interactions reported for breast cancer cells during matrix-deprivation (**Fig 2A**): a) AMPK and Akt can switch back and forth between their phosphorylated (active) and dephosphorylated (inactive) forms, b) phosphorylated AMPK (pAMPK) can upregulate the levels of PHLPP2 which promote the dephosphorylation of Akt, and c) phosphorylated Akt (pAkt) can upregulate the levels of PP2Cα associated with AMPK, thus enhancing AMPK dephosphorylation. These interactions are represented via a set of four coupled ordinary differential equations (ODEs). Each ODE tracks temporal evolution of the levels of AMPK, Akt, PHLPP2, PP2Cα, and the set of ODEs is solved numerically to obtain the steady state values for each of these four variables. To identify the robust dynamical features of this set of experimentally identified interactions, the kinetic parameters were chosen from a biologically relevant range of values (**Table S1**; see Materials and Methods). We chose 10,000 such unique parameter sets to represent the effects of cell-to-cell heterogeneity and 1000 initial conditions for each parameter set to characterize all the possible phenotypes across parameter sets.

**Figure 2:**
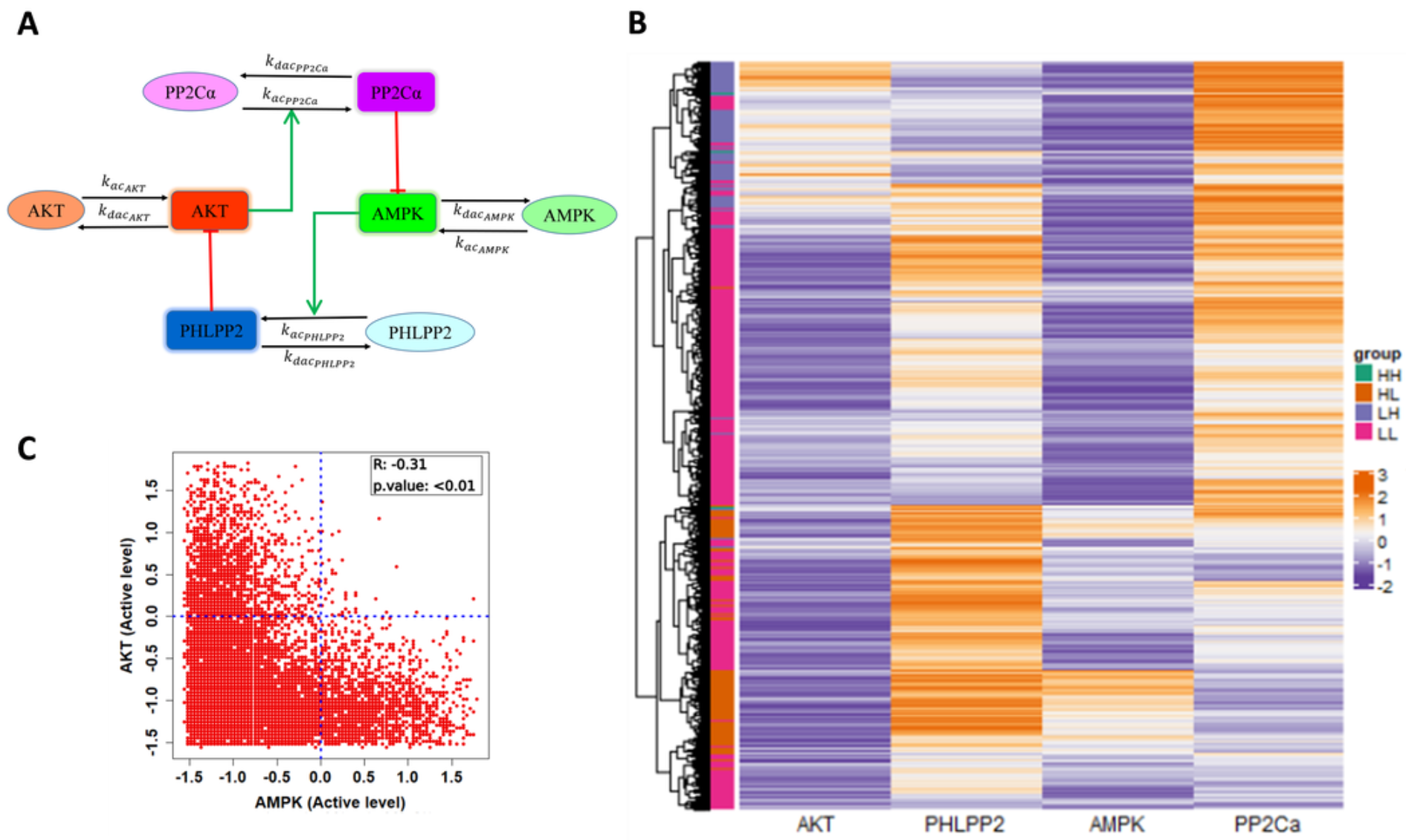
Dynamics of AMPK-Akt feedback loop. **A)** Reduced network considered in the model containing a total of 8 species – each of the four molecules (AMPK, Akt, PHLPP2, PP2Cα) has an active (rectangle) and an inactive (oval) form. Black arrows show the conversion of species from active to inactive and vice-versa. Red hammerheads represent inhibition, green arrows represent activation. Unless stated otherwise, further analysis shows the levels of only active species of the four molecules. **B)** Heatmap of steady states attained by 10,000 random parameter sets generated from 1000 random initial conditions of the active levels of AMPK, Akt, PHLPP2, PP2Cα. Color is based on the z-score calculated for the whole set of simulations, orange represents positive z-score (high) and purple represents negative z-score (low). LL, HL, LH and HH denote the four states when considering the steady state levels of pAMPK, pAKT - pAMPK^low^/ pAkt^low^, pAMPK^high^/pAkt^low^, pAMPK^low^/ pAkt^high^ and pAMPK^high^/pAkt^high^. **C)** Scatter plot of z-scores of AMPK and Akt steady state values represented in the heatmap showing the distribution of states. Pearson correlation coefficient, p-value are reported.

We collated the levels of AMPK, Akt, PHLPP2 and PP2Cα obtained from all parameter combinations and plotted them using a heatmap. We observed the emergence of four major clusters – pAMPK^low^/ pAkt^low^, pAMPK^high^/pAkt^low^, pAMPK^low^/pAkt^high^ and pAMPK^high^/pAkt^high^ state with pAMPK^high^/ pAkt^high^ state being the least frequent and pAMPK^low^/ pAkt^low^ state being the most frequent (**Fig 2B, S1**). A scatter plot between the pAMPK and pAKT levels revealed a significantly negative correlation (**Fig 2C, S1**), suggesting that pAMPK^high^/pAkt^low^ and pAMPK^low^/pAkt^high^ states can be dominant outputs of the network. The other nodes of the network also showed expected correlations trends (**Table S2**). However, the model suggests possible existence of pAMPK^low^/pAkt^low^ and pAMPK^high^/ pAkt^high^ states as well; the biological evidence for their existence yet remains inconclusive. The trend of predominance of pAMPK^high^/pAkt^low^ and pAMPK^low^/pAkt^high^ states was further confirmed by observations that the high frequency of pAMPK^low^/ pAkt^low^ was a consequence of parameter sampling (**Fig S2, Table S4**), i.e. in cases where the net activation rate of AMPK or Akt was quite low, independent of the AMPK or Akt activity (i.e. effects of PHLPP2 or PP2Cα). Therefore, we focused our attention to the pAMPK^high^/ pAkt^low^ and pAMPK^low^/pAkt^high^ states, and the parameter sets that converged to them. In particular, we focused on parameter sets that enabled a co-existence of these two states (bistability).

### The two states (pAkt^high^/ pAMPK^low^ and pAMPK^high^/pAkt^low^) can co-exist and stochastically switch between one another

To better understand bistability in this system, we performed nullcline and bifurcation analysis on the parameter sets showing bistability. First, for each parameter set, we constructed nullclines for pAMPK and pAKT. For a two-component system such as this, a nullcline represents the steady state levels of one component obtained over a range of values of the second component with its (i.e. the second component’s) rate of change over time being set to zero. The intersections of nullclines are, therefore, the points at which both the rate of change for both the components is zero, i.e., a steady state of the system. The advantage of this approach over dynamical ODE-based simulations is that we can identify unstable steady states as well, that act as tipping points for transition from one stable steady state to another. In other words, for these parameter states, the two identified stable states can co-exist and can transition from one to another.

Here, for a given parameter set, we first calculated the steady state levels of active Akt (pAKT) for different constant values of active AMPK (pAMPK) (red curve in **Fig 3A**, i). Next, we calculated the steady state levels of pAMPK for different fixed levels of pAkt (green curve in **Fig 3A**, i). These two curves are called as nullclines and their intersections identify the steady states of the system, two of which are stable (filled circles), and one unstable (hollow circle). The two stable states are pAkt^high^/ pAMPK^low^ and pAMPK^high^/pAkt^low^. The unstable state acts as a ‘tipping point’ beyond which a perturbation can allow switch from one state to another. Similar dynamical behavior was seen for other parametric combinations too, suggesting underlying bistability of the AMPK-Akt feedback loop (**Fig 3A**, ii-iii).

**Figure 3:**
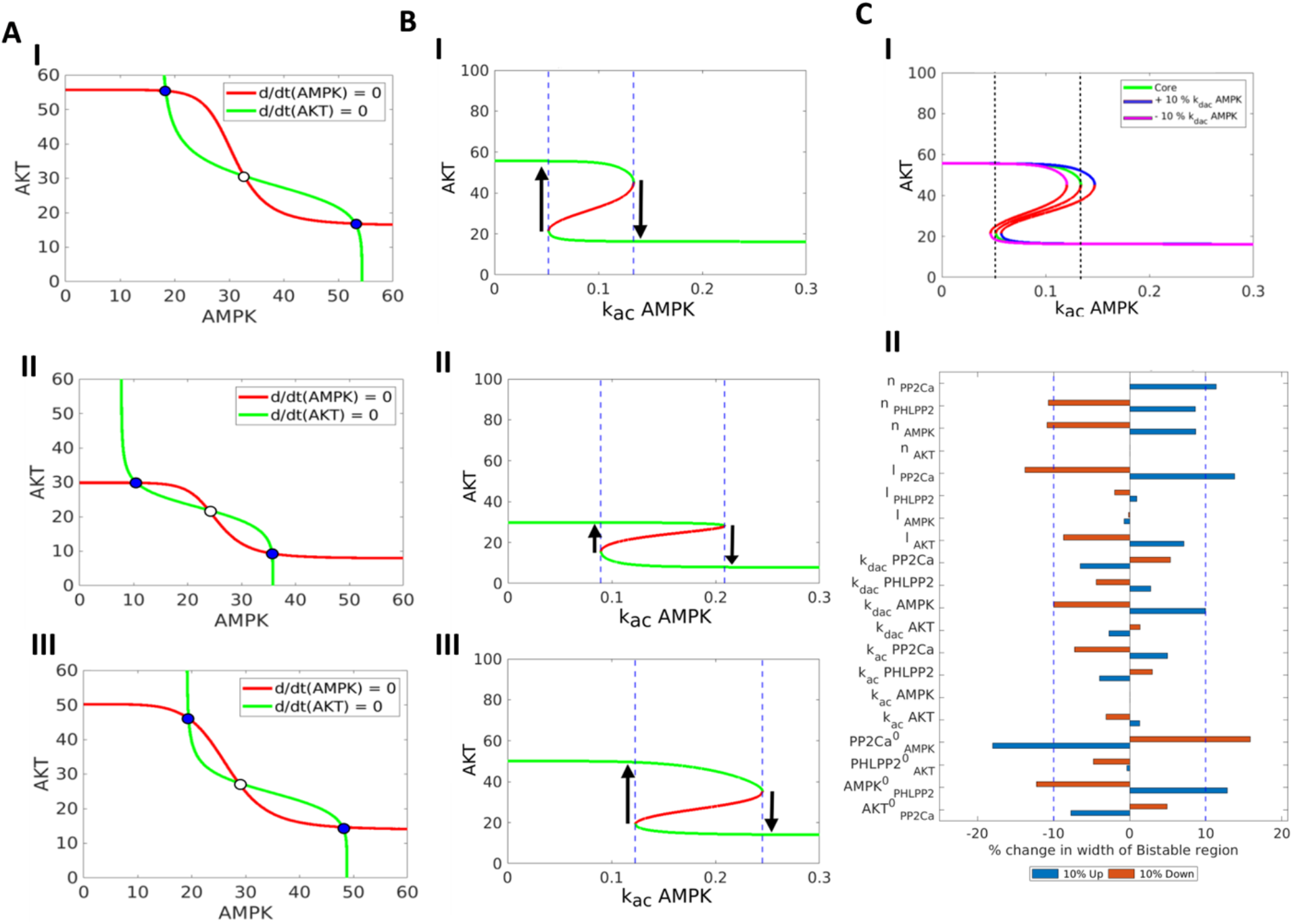
Nullcline, bifurcation and sensitivity analysis. **A)** Nullclines for three representative bistable parameter sets (rows 1-3 in **Table S3**). Red curve is AMPK Nullcline [d/dt(AMPK)=0]. Green curve is Akt Nullcline [d/dt (AKT)=0]. Blue circles represent stable states, white circles represent unstable steady state. **B)** Bifurcation of Akt steady state levels with respect to the activation rate of AMPK (k_ac AMPK) for three representative bistable parameter sets (rows 1-3 in **Table S3**).Green curves denote stable states and red curves denote unstable states. Region bound by the blue dashed line represents the bistable region, where both states can co-exist. **C)** i) Representative bifurcation of Akt levels with respect to activation rate of AMPK (k_ac AMPK), drawn for three different levels (control, ±10% change in deactivation rate of AMPK (k_dac AMPK). Green curve shows the control case, blue curve for + 10 % and magenta Curve for – 10 % of k_dac AMPK. ii) Sensitivity analysis of the width of the bistable region (length of the segment of the x axis between the black dotted lines in bifurcation diagram) for a parameter set (row #1 in **Table S3**) with changes in individual parameter by ±10% from the original value. Blue dotted line represents the ±10 % change. Arrows show possible transitions between the different states.

To investigate how this feature of bistability may depend on various kinetic parameters in the model, we plotted a bifurcation diagram that tracks the levels of pAKT at varied values of activation rate of AMPK (k_ac_AMPK). We chose k_ac_AMPK as the bifurcation parameter to reflect the cases where the activation rate of AMPK can be altered by cell-intrinsic or cell-extrinsic factors. At low k_ac_AMPK values (< 0.05), pAkt levels are relatively higher, because the dephosphorylation of Akt by AMPK is weaker. Similarly, at high K_ac_AMPK values (> 0.13), pAkt levels are relatively lower, because of strong inactivation of Akt by AMPK. However, at intermediate range of these values (area bounded between dotted blue lines), a cell can exhibit bistability in terms of pAMPK levels, i.e. it can exist in either a pAkt^high^/ pAMPK^low^ or pAMPK^high^/pAkt^low^ state (**Fig 3B**, i). Similar trends are seen for other parameter sets shown earlier (**Fig 3B**, ii-iii).

To quantify the range of bistability in the bifurcation diagram and its dependence on other kinetic parameters besides k_ac_AMPK, we plotted this bifurcation at different values of activation rate of Akt (k_ac_Akt) and observed that these curves (blue and magenta curves in **Fig 3C**, i) largely overlapped with that seen earlier (green curve in **Fig 3C**, i). Next, we varied each parameter, one at a time, by +/- 10% and calculated the percentage change in the range of k_ac_AMPK values enabling bistability, i.e. the distance between the dotted vertical lines. This sensitivity analysis suggested that the percentage change in this range was above 10% for only a few parameters such as those defining the effect of PP2Ca on AMPK (**Fig 3C**, ii; **Fig S3, S4**). These results underscore that the bistability feature of AMPK-Akt feedback loop is largely robust to parameter variations, and consequently, cells in an isogenic population may exist in two distinct states: pAkt^high^/ pAMPK^low^ and pAMPK^high^/pAkt^low^.

Next, we varied two parameters simultaneously, to map the two-dimensional parameter region in terms of (co-)existence of the two states. At high K_ac_AMPK and low K_ac_Akt, only the pAMPK^high^/pAkt^low^ state was observed. Similarly, low K_ac_AMPK and high K_ac_Akt allowed only the pAkt^high^/ pAMPK^low^ state. Both these states co-existed in a parameter region lying between these extremes (yellow region in **Fig 4A**, i). These trends were reinforced based on bifurcation diagrams drawn for an intermediate value of k_ac_Akt (= 0.15), with k_ac_AMPK as the bifurcation parameter. Bistability existed for intermediate values of k_ac_AMPK; while higher or lower values led to monostable regions where only one state existed (**Fig 4A**, ii). Bifurcation diagram with k_ac_Akt as the parameter confirmed the trend (**Fig 4A**, iii). Similar characteristics dynamics was seen for other parameter sets (**Fig S5).**

**Figure 4:**
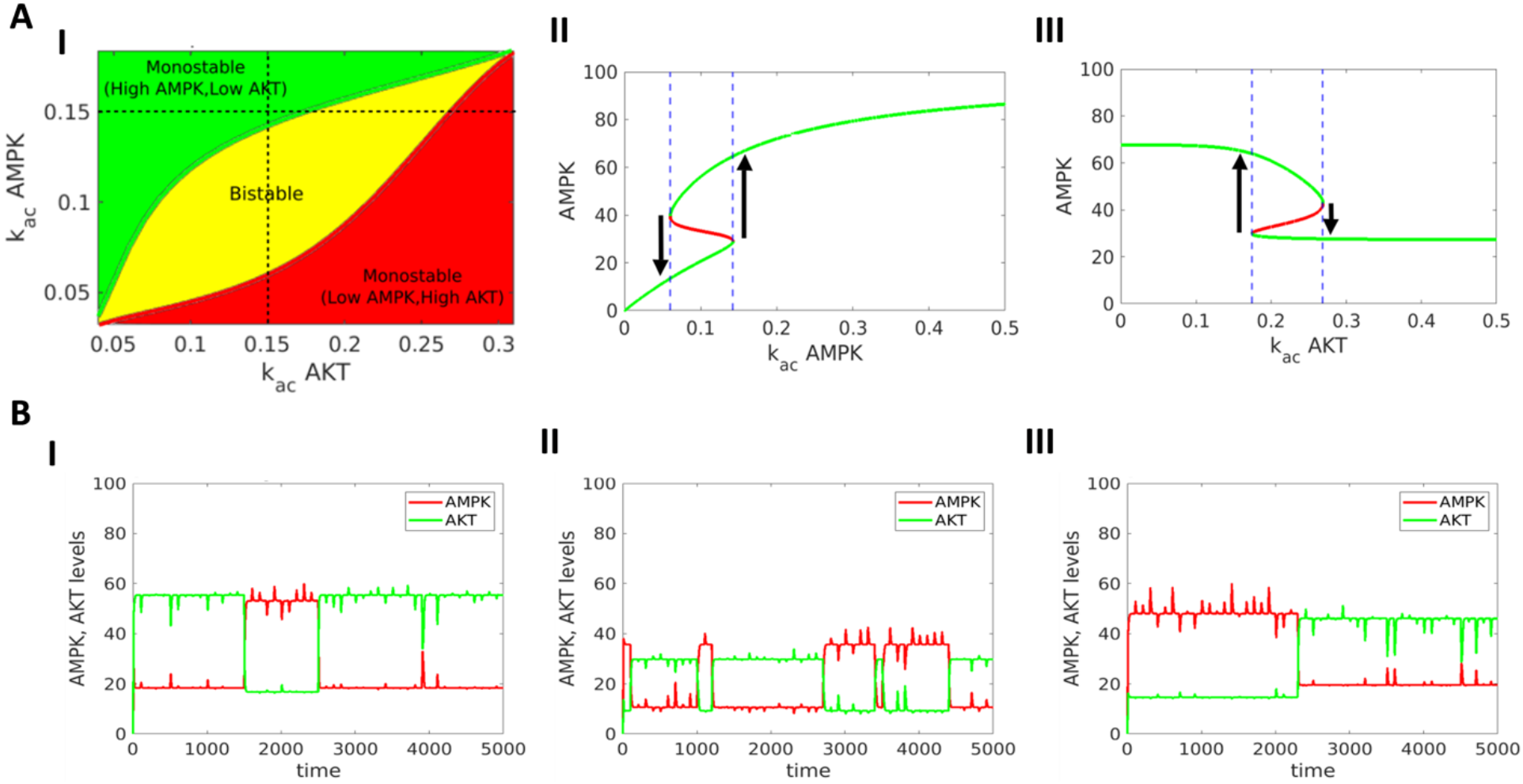
Phase plots and stochastic simulations. **A)** i) Phase diagram for two parameters - activation rates of AMPK and Akt - showing monostable and bistable regions. All other parameter values correspond to those given in row #1 in Table S3. ii) Bifurcation diagram of AMPK levels with respect to k_ac AMPK for a constant value of k_ac Akt = 0.15. iii) Same as ii) but with respect to k_ac Akt for a constant value of k_ac AMPK=0.15. Green curve shows stable states, red curve shows unstable states. Blue dotted lines show region of bistability. **B)** Stochastic simulations showing trajectories of AMPK, Akt values under the influence of noise for three representative parameter sets (rows 1-3 in **Table S3**). Noise parameter value η = 20.

The co-existence of two (or more) states in a bifurcation diagram indicates that they may ‘spontaneous’ switch among one another. Thus, we performed stochastic simulations to examine for state switching under the influence of biological noise. These simulations revealed that cells can switch back and forth between the pAkt^high^/ pAMPK^low^ and pAMPK^high^/pAkt^low^ states for varying strengths of noise parameter (**Fig 4B, S6**), highlighting that spontaneous state switching may be an outcome of the AMPK-Akt loop.

Such state switching implies that when a cell population is sorted into subpopulations, it is possible for a subpopulation to give rise to another and potentially generate a population distribution similar to that seen in parental population [30,31]. The rates of switching to and from a subpopulation to/from another one may be unequal, depending on the relative stability of the two (or more) states, evidenced by different mean residence times in a given state [32].

### Experimental and clinical data supports the model predictions of bistability in AMPK-Akt loop

To experimentally interrogate our observations of spontaneous state switching between pAkt^high^/ pAMPK^low^ and pAMPK^high^/pAkt^low^ states, we performed FACS based cell sorting experiments in MDA-MB-231 cells stable for EGR1(promoter)-TurboRFP (EGR1 promoter-reporter system). In this approach, EGR1 promoter is used as a readout for AMPK activity, where high EGR activity (assessed by RFP intensity) corresponds to low AMPK activity and *vice-versa* [33]. Cells were grown in attached condition and sorted into high and low RFP group by FACS. These cells were cultured again for the indicated times in attached condition and we observed that these cells regained their original heterogeneous nature with time (**Fig 5A**).

**Figure 5:**
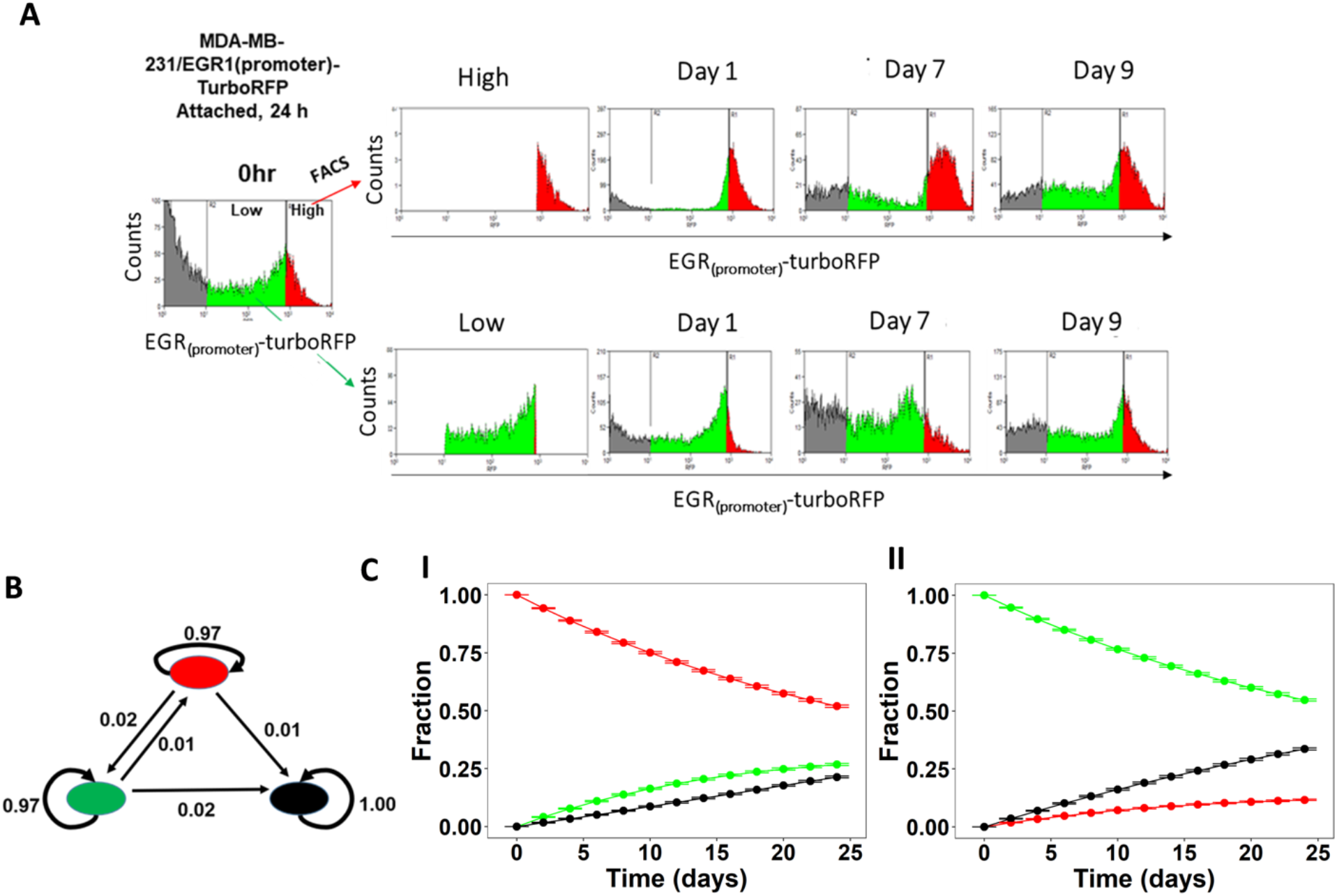
Stochastic state transitions. **A)** Experimental validation of AMPK-Akt feedback loop using MDA-MB-231 EGR1-Turbo RFP cell lines sorted for high and low RFP expressing population using FACS. The RFP high population (red) corresponds to low pAMPK and high pAkt, and RFP^low^ (green) population corresponds to low EGR activity, high pAMPK, and low pAkt. Grey population correspond to cells that have lost the vector. Histograms show the population composition after 1,7 and 9 days when started with distinct RFP^high^ (red, top panel) and RFP^low^ (green, bottom panel) populations. **B)** State transition graph: each node is a cell phenotype colored the same as FACS data for representative purposes, each edge represents transition between the corresponding phenotypes. Transition rates (per day) calculated from the Markov model are shown on the corresponding arrows. **C)** Predicted evolution of population heterogeneity when started from the initial population fractions equivalent to distinct populations in the FACS data, using the transition probabilities calculated here. Error bars represent mean ± standard deviation from 1000 such simulations.

To quantify the observed phenotypic transitions between low and high AMPK (red and green populations respectively) cells, we constructed a discrete time Markov chain model [31]. The model assumes that transition rates between these cell types are independent of each other and do not change with time. We further assumed that the non-expressing cells (black population in the FACS distributions) do not transition into any other cell type. Using the R package CellTrans [34], we obtained the transition matrix, an ordered collection of transition rates (**Fig 5B**). Using these transition rates, we simulated the evolution of population composition starting from homogeneous populations (**Fig 5C**). The model predicts a higher transition rate from pAkt^high^/pAMPK^low^ to pAMPK^high^/pAkt^low^ phenotype, than that of pAMPK^high^/pAkt^low^ to pAkt^high^/ pAMPK^low^, suggesting that the pAMPK^high^/pAkt^low^ population may be relatively more stable.

Further, to support our finding in the clinical setting, we performed a correlation of pAkt and pAMPK levels in patient samples using RPPA data from the TCGA cohort. A significant negative correlation was observed between pAkt and pAMPK levels across various cancer types (**Fig 6, Table S5**). These data further support the existence of bistability in the dynamics of AMPK-Akt feedback loop. Put together, our integrated computational-experiment analysis shows that the AMPK-Akt crosstalk can give rise to bistability and can lead to switching of states, driving heterogeneity at population level.

**Figure 6:**
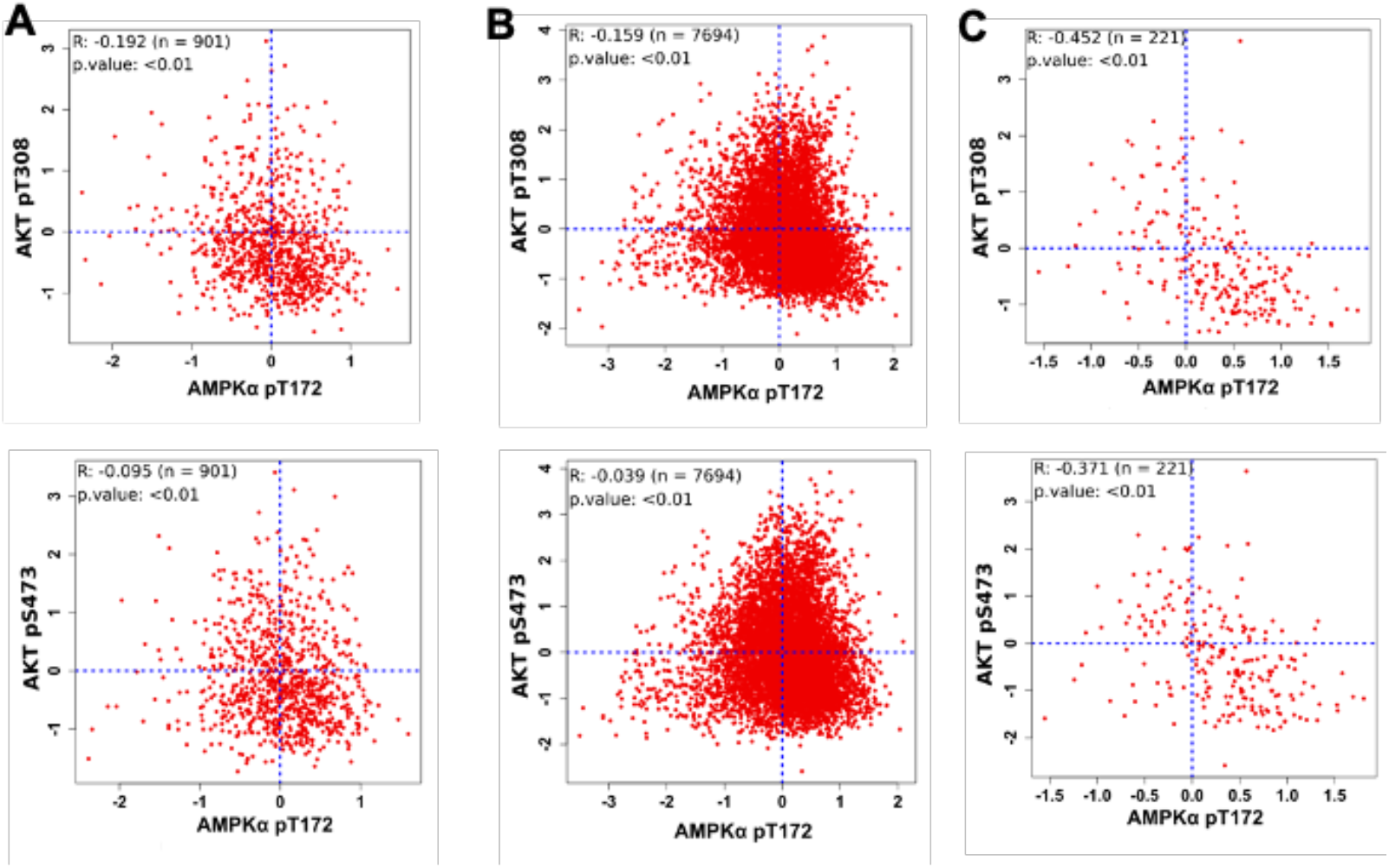
Clinical validation of AMPK-Akt double negative feedback loop. Scatter plot of active levels of AMPK (AMPK pT172) and Akt (pT308 and pS473) in **A)** Breast cancer cohort (TCGA-BRCA)[n=901], **B)** Pan Cancer cohort of 32 cancer types [n=7694] and **C)** Sarcoma cohort (TCGA-SARC) [n=221]. Each dot represents one patient and coordinates correspond to the protein expression levels capture using RPPA (reverse phase protein array) with Akt and AMPK active forms (AMPK - pT172, Akt-pT308 and Akt-pS473). Pearson’s correlation coefficients and p-value are reported.

## Discussion

Phenotypic switching can play crucial roles during cancer progression, as seen across cancer types. Prostate cancer cells can switch to a neuroendocrine-like state that is refractory to various therapies [35]. Similarly, in small cell lung cancer, cells under therapy-induced stress can reversibly switch to a hybrid neuroendocrine/mesenchymal state [36]. Besides these specific examples, EMP and CSCs are archetypal examples of phenotypic switching reported in many carcinomas [37,38] and non-epithelial tumors too [39-42]. Recent progress in collecting high-throughput spatiotemporal data and mapping the regulatory networks underlying these axes of plasticity has led to developing complex mechanism-based and population-based mathematical models to decode dynamical traits of phenotypic switching such as dose-dependence, reversibility, hysteresis and transition rates among the cell states [43-51]. A hallmark of regulatory networks enabling phenotypic switching in cancer cell populations is multistability, i.e. the ability of isogenic cells to reversibly acquire diverse phenotypes [5,11,12,52-56], as reported earlier for bacterial [57] and viral [58] populations too. Multistability can enable ‘spontaneous’ switching among cell phenotypes (different attractors in the Waddington’s landscape) due to biological noise (that can operate at multiple levels including transcriptional or conformational [59,60]), and thus facilitate non-genetic heterogeneity [8,61]. For instance, in the context of EMP, PMC42-LA cells showed a bimodal distribution of EpCAM, and either subpopulation (EpCAM-^high^ or EpCAM-^low^) was capable of generating the other without any exogenous overt induction [48]. Similar observations have been reported in maintaining a dynamic equilibrium of CSCs and non-CSCs in breast cancer [62].

Here, we demonstrate breast cancer cells switching between pAkt^high^/pAMPK^low^ and pAMPK^high^/pAkt^low^ states in adherent cell populations. Together with reinforcing observations for switching among these subpopulations in matrix-detached conditions too, using the same reporter construct in MDA-MB-231 cells [33], our results strongly support the existence of multistability in AMPK-Akt double negative feedback loop, as predicted by our mechanism-based mathematical model. A limitation of our mathematical model is the limited set of interactions between AMPK and Akt that we incorporated for the purpose of characterizing their dynamics in the context of matrix-deprivation. Other contextdependent interactions between AMPK and Akt have been reported, for instance, AMPK can activate Akt in acute lymphoblastic leukemia [63]. Also, AMPK mediated phosphorylation of Skp2 at S256 can activate the Skp2 SCF complex, driving K63-linked ubiquitination and eventual activation of Akt [64]. However, these interactions have not been yet observed in matrix-deprivation conditions. Such context-specific differences may allow for other dynamics for AMPK and Akt, such as oscillations seen in MCF10A cells upon inhibition of glycolysis and mitochondrial ATPase [65]. Intriguingly, AMPK can form double negative feedback loops with other molecules such as mTORC1 [66], which is involved in a similar feedback loop with ULK1 [67]. Coupling of such ‘toggle switches’ can influence the emergence of multistability [68].

Another salient feature of multistability is hysteresis, as observed for bacterial cells [69] and for EMP in cancer cells [11]. Future experiments should investigate the possibility and implications of hysteresis in AMPK-Akt feedback loop driving the adaptation of breast cancer cells to matrix-deprivation (in other words, anchorage-independence) stress. Also, anchorage-independence has been shown to be associated with other axes of phenotypic plasticity: EMP [70–72], CSCs [73], and metabolic reprogramming [74–76]. However, a systems-level understanding of coordination of cell states along these interconnected axes remains elusive. Future integrative computational-experimental efforts, similar to the approach taken here, can be critical in investigating such coupled dynamics of phenotypic plasticity during the challenging metastatic cascade and identify therapeutic targets that can impact multiple axes of plasticity simultaneously.

## Acknowledgements

This work was supported by Dr. Vijaya and Rajagopal Rao Biomedical Research Fund awarded to Centre for BioSystems Science and Engineering, Indian Institute of Science (IISc), Bangalore (MKJ). AR acknowledges support from DBT-IISc Partnership Program Phase-II (BT/PR27952-INF /22/212/2018), and support from DST-FIST and UGC, Govt. of India, to the department of MRDG, IISc. AR is a recipient of the Welcome Trust/DBT India Alliance Fellowship. AC acknowledges the Junior Research Fellowship (JRF) awarded by the Council of Scientific & Industrial Research (CSIR), India. KH was supported by Prime Ministers’ Research Fellowship (PMRF) awarded by the Ministry of Human Resources and Development (MHRD), Government of India. Authors acknowledge MRDG and IISc FACS facility.

## Author contributions

MKJ conceived research; MKJ and AR supervised research; AC, KH and SK performed research; all authors analyzed data and participated in writing and editing of the manuscript.

## Materials and methods

### ODE model of the AMPK-Akt network

The dynamics of the species in the regulatory network (**Fig 2A**) are represented using a system of Ordinary differential equations (ODEs) given below:

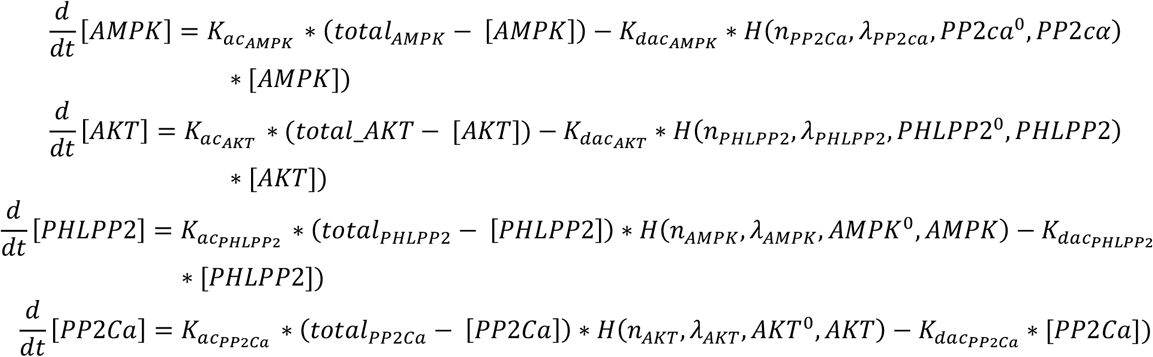

Here, each equation represents the rate of change of the levels of the entity given in the LHS of the equation. The activation and deactivation rates of the species are given as *K_acx_* and *K_dacx_*, (*X* ∈ {*AMPK, AKT, PHLPP2, PPCA*}) respectively. The regulatory interactions are represented using shifted hill function form [77] that takes the form given below

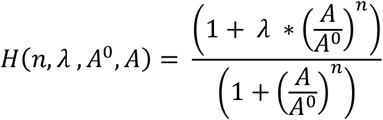

Where A is the effector species, n is the Hill’s coefficient that represents the non-linearity of the regulatory interaction, *λ* is the maximum fold change in activation/deactivation rate caused by A. *λ*> 1 indicates activation and *λ*<1 indicates inhibition, A^0^ is the Activation/Inhibition threshold level of A. The parameter value ranges with the corresponding dimensions are given in Table S1.

### Temporal Profiles and Steady state estimation

10000 Parameter sets were randomly sampled from the ranges mentioned in Table S1. For each parameter set, the ODE system was simulated using 1000 randomly generated initial conditions between the range [0, total molecules]. The temporal profiles for each initial condition were computed numerically using the *ode23* function of Matlab *R2019A* until steady state was reached for all species or 1000 time-steps, whichever was smaller.

### Nullcline, Bifurcation and Phase plane analysis

For a given parameter set for the ODE system, nullclines for AMPK and Akt were obtained by setting their respective rates of change to zero and using constant value for the level of that species. The corresponding steady state of the other variable is obtained by temporal simulations of the modified ODE system as described above. Bifurcation analysis was done using the software package MATCONT [78]. Phase planes were obtained by combining multiple bifurcations generated k_ac_AMPK within the range given the Table S1.

### Noise induction

We used a simple formalism to induce noise that introduces a white noise sampled from a normal random distribution of mean 0 and variance *σ, σ* ∈ {20,30,40}. This noise is added to the species levels at fixed intervals in the simulation, representing the additive stochasticity to the species levels in the time period of simulation.

### Clinical Data

Reverse Phase Protein Array (RPPA) dataset for TCGA-Pan Cancer 32, Breast Cancer and Sarcoma were downloaded from https://www.tcpaportal.org/tcpa/download.html. Pearson’s correlation analysis between phosphorylated AMPK (pT172) and phosphorylated AKT (pT308 and pS473) levels were calculated using cor.test() function from stats package and Scatterplot was generated using plot() function in R 3.6.1.

### Cell line and culture condition, FACS sorting and analysis of the plasticity

Breast cancer cell line MDA-MB-231 was procured from ATCC and validated subsequently by STR analysis. These cells were cultured in DMEM (Sigma-Aldrich) supplemented with 10% FBS containing penicillin and streptomycin, at 37°C in 5% CO2 incubator. Cells were trypsinized and counted before seeding for every experiment. For FACS sorting purposes, MDA-MB-231 cells stably expressing EGR1promoter-TurboRFP (readout for the ERK activity) were sorted into high and low RFP cells after culturing them on 90 mm tissue culture dish. These sorted cells were cultured in attached condition for the indicated time-points. Analysis of the high and low RFP sorted cells for their ability to attend the original heterogeneity was performed after the indicated time point of the culture in attached condition. Representative data were analysed using Summit software V5.2.1.12465.

### Markov chain modelling and simulations

For calculating the transition matrices for the discrete time markov chain (DTMC) model, we analysed the FACS data using the R package CellTrans. Briefly, the package makes use of the fact that the phenotypic frequency distribution at time t in a DTMC is obtained by multiplying the initial distribution with the transition matrix raised to the power of t and calculates the transition matrix from the NxN phenotypic distribution matrix (PDM) at time t and that at the beginning of the experiment. Typically, the initial populations are FACS sorted, hence the initial PDM is an NxN identity matrix. Using transition matrices, we simulated the trajectories of sorted cell populations. In an instance of the simulations, we start with a sorted population of size 10000. At each time step, the transition of a given cell from its current phenotype to a new phenotype is decided using a uniform random number and the row of the transition matrix corresponding to the current phenotype of the cell. 1000 simulations were performed to obtain a distribution of the population fractions of cell phenotypes at each time point.

### Code availability

The corresponding codes are available at https://github.com/csbBSSE/AmpkAkt.

## Supplementary Table Legends

**Table S1:** Sheet 1 shows the literature reported values of parameter values. Sheet 2 shows the parameter ranges used for generating parameter sets.

**Table S2:** Shows the Pearson’s correlation values and corresponding p values for the comparisons between AMPK, Akt, PHLPP2 and PP2Cα for three replicates.

**Table S3:** Shows the 10 representative parameter sets used in the study.

**Table S4:** Shows the mean and standard deviation of k_ac/k_dac AMPK and Akt values for four groups (LL, HL, LH and HH), and p values for pairwise comparisons between the groups.

**Table S5:** Shows the Pearson’s correlation values and p values between AMPK (pT172) and Akt (pT308 and pS473) for 32 different TCGA cancer cohorts.

