## Supplementary material for "Multistability and consequent phenotypic plasticity in AMPK-Akt double negative feedback loop in cancer cells": SI Tables, Figures: Supplementary_figures_final.pdf

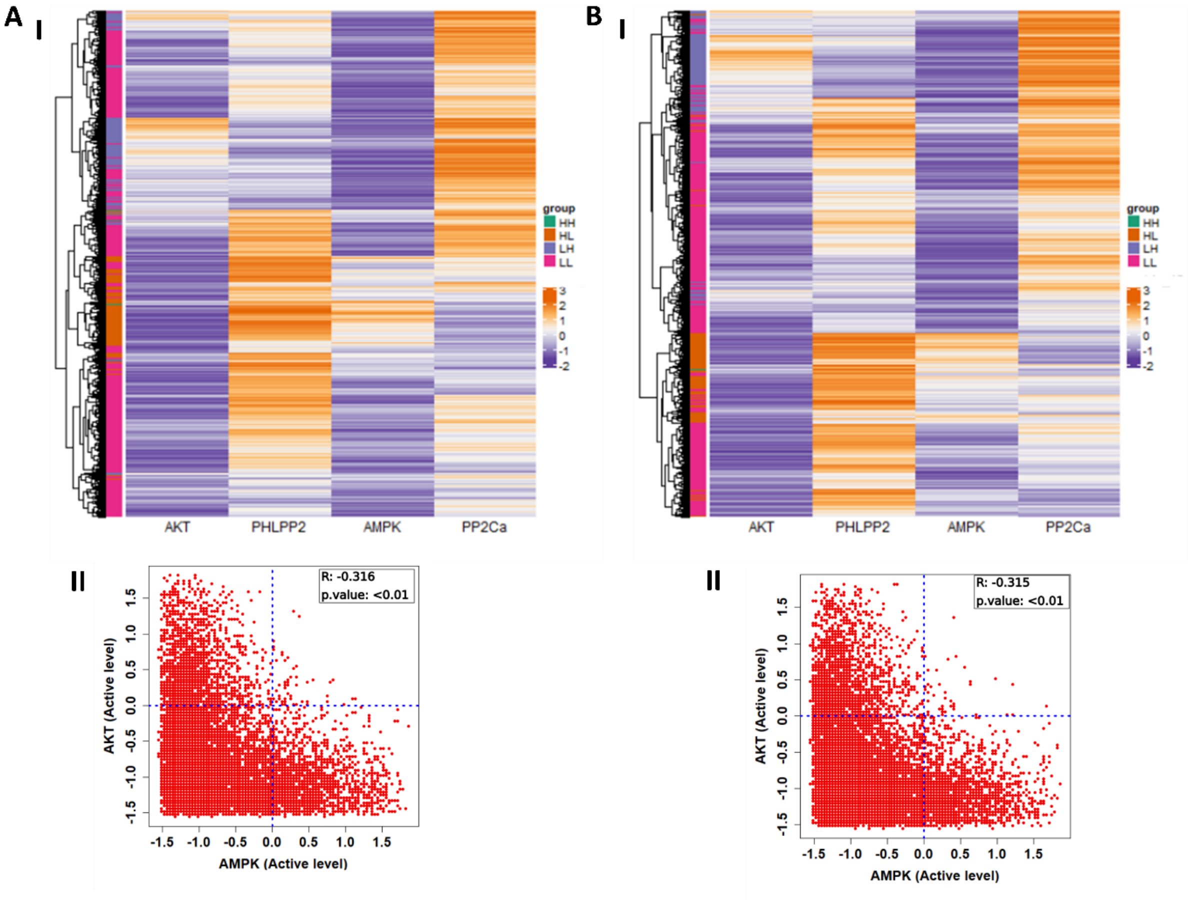


**Figure S1:**

**A-B (I)** Heatmap of final states attained by 10,000 random parameter sets generated from 1000 random initial states of the active levels of AMPK, AKT, PHLPP2, PP2Cα. Color range of the cells is based on z-score calculated for the whole set, orange represents positive z-score, and purple represents negative z-score. LL, HL, LH and HH denote the four states - $pAMPK^{low}$/ $pAkt^{low}$, $pAMPK^{high}$/$pAkt^{low}$, $pAMPK^{low}$/ $pAkt^{high}$ and $pAMPK^{high}$/$pAkt^{high}$.

**(II)** Scatter plot of AMPK and AKT z-score values represented in the heatmap emphasising the distribution of states. Pearson correlation coefficient, p-value are reported.

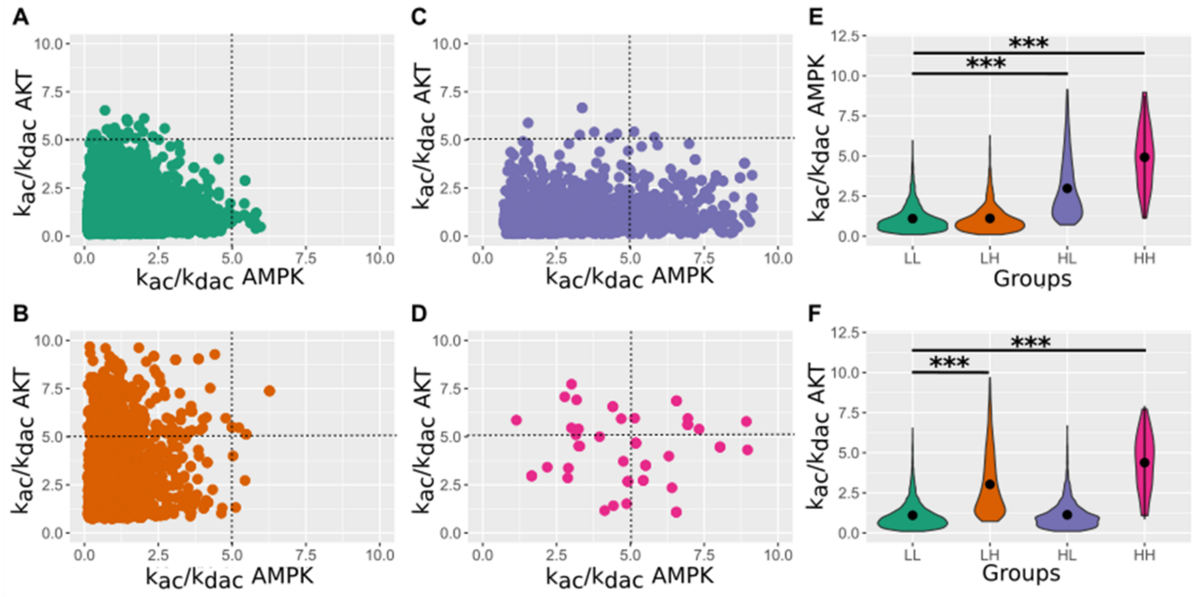


**Figure S2:**
**A-D** Scatter plot for k_ac/k_dac AMPK and Akt for four groups (LL, HL, LH and HH). **E, F** Violin plots showing the distribution k_ac/k_dac AMPK and Akt values across different groups. Black dot represents the mean of the distribution and *** denotes p-value <10e-5. LL, HL, LH and HH denote four states respectively
- pAMPK$^{low}$/ pAkt$^{low}$, pAMPK$^{high}$/pAkt$^{low}$, pAMPK$^{low}$/ pAkt$^{high}$ and pAMPK$^{high}$/pAkt$^{high}$.

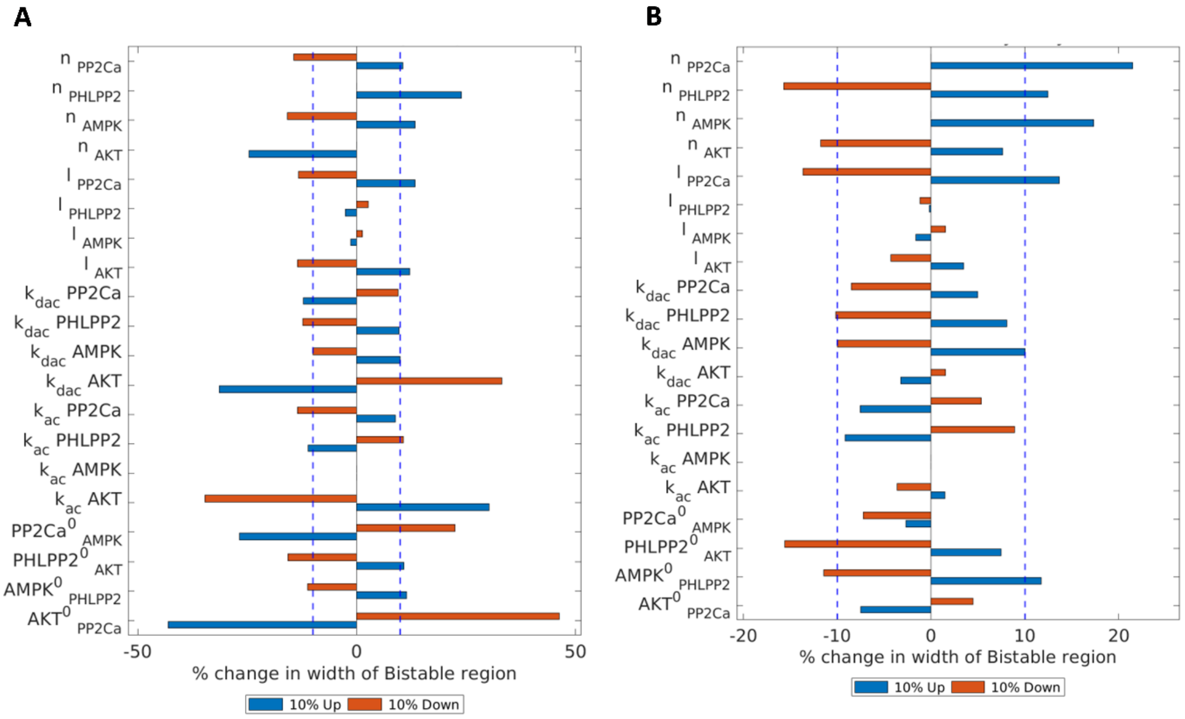


**Figure S3:**
Sensitivity of the width of the bistable region for a parameter set with changes in individual parameter by ±10% from the original value. Blue dotted line represents the ±10 % change. A, B show the results for parameter set 2-3 (rows #2-3 in Table S3) respectively.

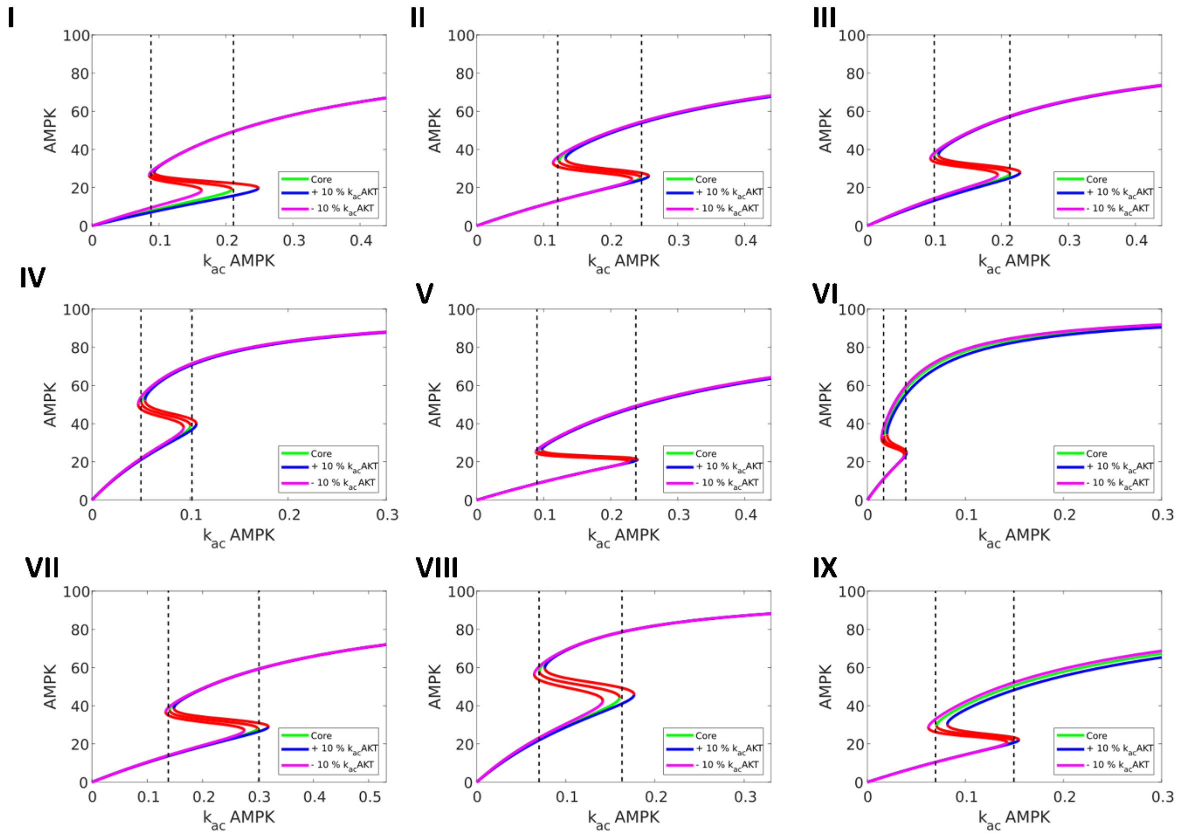


**Figure S4:**
Representative bifurcation of AMPK levels with respect to (±10% change) in the activation rate of AMPK (k_ac AMPK). Green curve is for the core value, blue curve for + 10 % and magenta curve for – 10 % of the activation rate of Akt (k_ac Akt). I – IX are for parameter sets 2-10 (row #2-10 in Table S3).

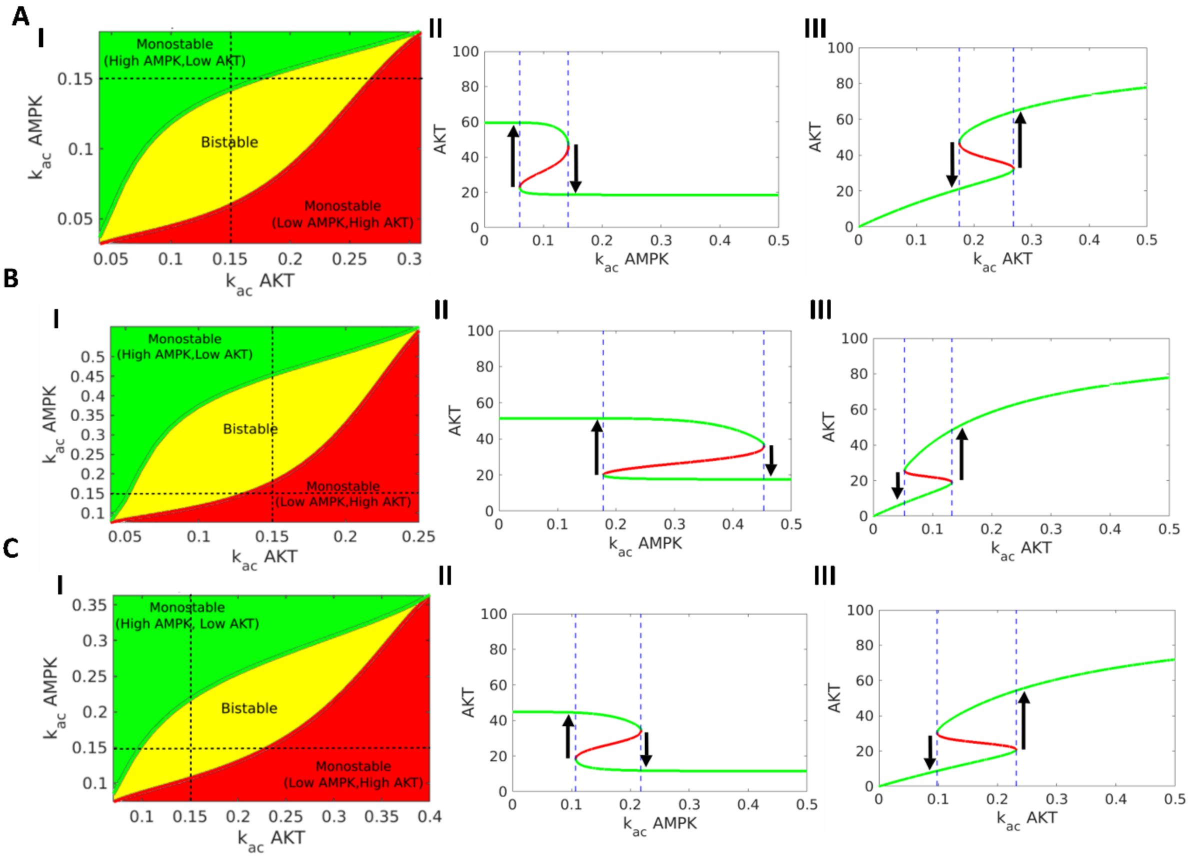


**Figure S5:**

(I) Phase diagram for two parameters –activation rates of AMPK and Akt – showing monostable and bistable regions. (II, III) Bifurcation of Akt levels with respect to k_ac Akt and k_ac AMPK under constant value (0.15) of the other parameter (k_ac AMPK and k_ac Akt) respectively. Green curve shows stable states, red curve shows unstable states. Blue dotted lines show region of bistability. A- C for parameter set in rows #1-3 in Table S3.

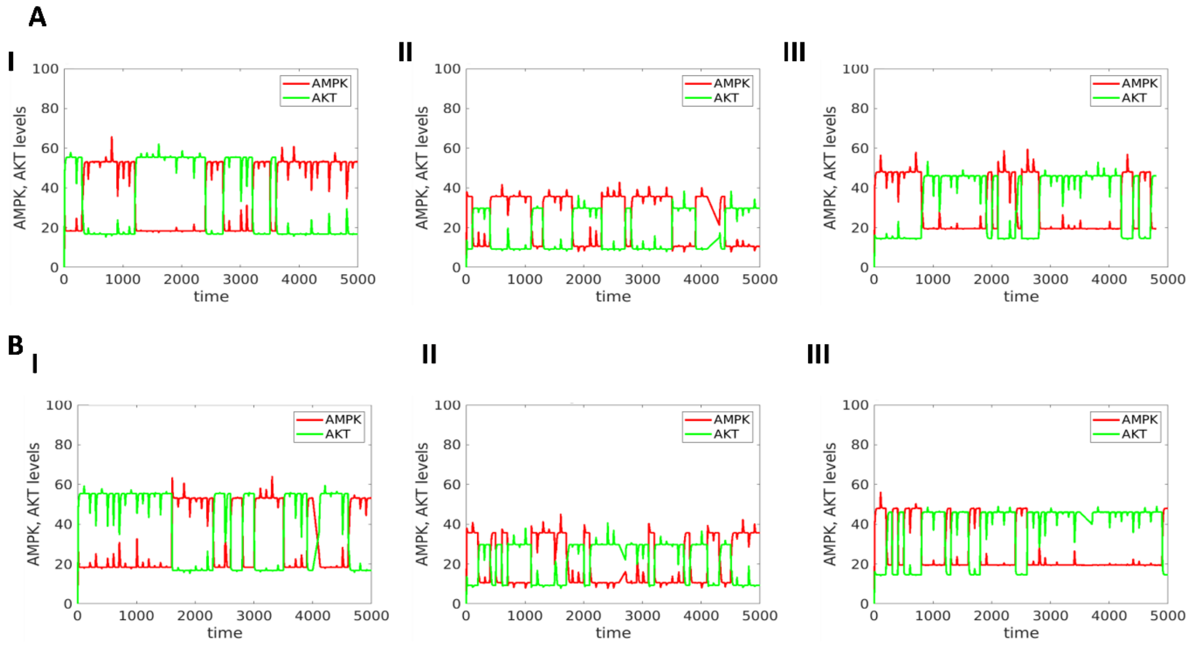


**Figure S6:**
Stochastic simulations showing trajectories of AMPK, Akt values under the influence of noise parameter **A** (η=30) and **B** (η =40) for three representative parameter sets (rows #1-3 in Table S3).

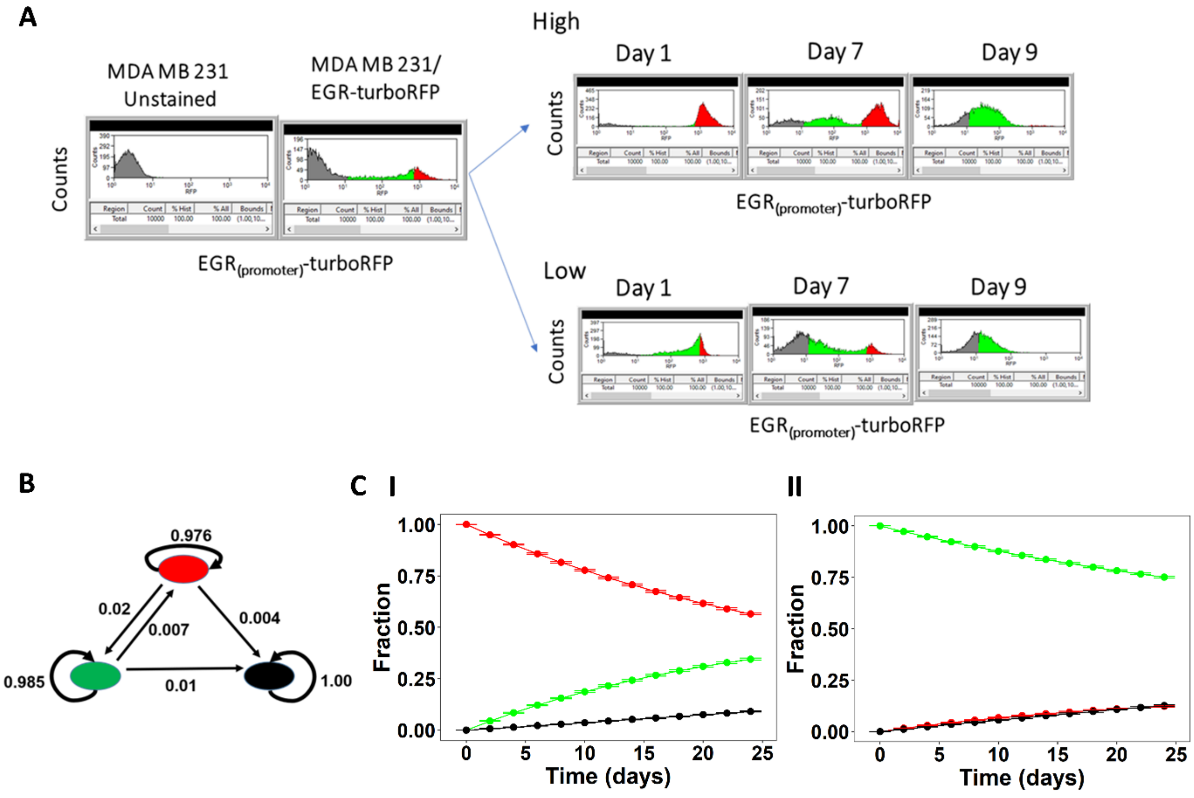


**Figure S7:** Same as Fig 5 but for another replicate.
